# Equivalence of the Transition Heat Capacities of Proteins and DNA

**DOI:** 10.1101/2021.12.29.474479

**Authors:** Matthew W. Eskew, Albert S. Benight

## Abstract

It has been reported for many globular proteins that the native heat capacity at 25°C, *per* gram, is the same. This has been interpreted to indicate that heat capacity is a fundamental property of native proteins that provides important information on molecular structure and stability. Heat capacities for both proteins and DNA has been suggested to be related to universal effects of hydration/solvation on native structures. Here we report on results from thermal denaturation analysis of two well-known proteins, human serum albumin and lysozyme, and a short DNA hairpin. The transition heat capacities at the T_m_ for the three molecules were quantitatively evaluated by differential scanning calorimetry. When normalized *per gram* rather than *per mol* the transition heat capacities were found to be precisely equivalent. This observation for the transition heat capacities of the proteins is consistent with previous reports. However, an identical transition heat capacity for DNA has not been reported and was unexpected. Further analysis of the collected data suggested a mass dependence of hydration effects on thermal denaturation that is preserved at the individual protein amino acid and DNA base levels. Equivalence of transition heat capacities suggests the possibility of a universal role of hydration effects on the thermal stability of both proteins and DNA.

## Introduction

For most globular proteins, it is well established that the native heat capacity at 25°C, p*er gram*, is universally the same ^1-4^. In this study, transition heat capacities *per gram* (heat capacity at the transition temperature, T_m_), were evaluated for two proteins and a DNA hairpin, and found to be equivalent. To our knowledge this equivalence for DNA and proteins has not been previously reported and implies a common linkage, in a thermodynamic sense, between their global structural stabilities.

In a conventional differential scanning calorimetry (DSC) experiment, the excess heat capacity C_p_ is measured as a function of increasing temperature. Plots of C_p_ versus T are called DSC thermograms ^5^. On thermograms of proteins and DNA, ΔC_p_ is the difference in heat capacity between the native (low temperature) and denatured (high temperature) states ^6-8^. ΔC_p_ indicates the relatively higher heat capacity of an unfolded molecule compared to its native state ^6-8^. ΔC_p_, *per mole*, has been found to be greater for proteins than DNA and attributed to differences in solvent exposed surface areas of denatured versus native states ^7-10^. Also obtained from DSC measurements is the absolute heat capacity corresponding to the maximum peak height on the thermogram at T_m_. The transition enthalpy and entropy are derived from integration of the thermogram. Differences in absolute heat capacities among globular proteins are due to differences in their partial specific volumes (psv) ^4^. The transition heat capacity is derived from the absolute heat capacity at T_m_ corrected for differences in psv for different proteins.

The major contribution to the absolute heat capacity for both proteins and DNA comes from water ^10-16^. Water molecules are essential for maintenance of native and active protein and DNA structures ^10-16^. In the process of denaturation of protein and DNA molecules, multiple interactions between water and the native and denatured states contribute significantly to the melting enthalpies and entropies ^7, 17^. Enthalpic and entropic contributions from interactions of water molecules with both polar and apolar amino acid residues in proteins have been extensively investigated ^10, 13^. Likewise, water interactions with the polar surface and hydrophobic core of duplex DNA have also been studied ^6, 9, 14, 18^. Disruption of native protein and DNA structures by thermal melting exposes both hydrophobic and hydrophilic regions; with disruption of the hydration shell of ordered water that accompanies melting of the overall structure ^12, 17, 19^. These hydration effects are thought to comprise the primary source of enthalpic changes in both proteins and DNA that accompany denaturation ^6, 7, 12-15, 17, 19^. Since calorimetric enthalpy is derived directly from excess calorimetric heat capacity, it stands to reason that hydration effects also represent the primary source of the absolute heat capacity.

If hydration effects are primarily responsible for the maximum calorimetric peak height, why are these values universal for proteins of vastly different sizes and structures? Our operating assumption is that, once normalized by mass, there are comparable amounts of solvent accessible area that are susceptible to the same hydration effects. Presumably, solvent accessible surface area scales linearly with mass.^13^ Then, for a sample containing a given mass of large macromolecules (proteins or DNA) there are relatively fewer individual molecules. In contrast, for samples containing a given mass of smaller macromolecules there will be relatively more individual molecules. Thus, at the same sample mass, the greater mass of large molecules is balanced by fewer molecules, while smaller molecules will have more molecules in the sample. For either case, this balance reveals, for a given mass, essentially equivalent solvent accessible areas available for solvation, regardless of molecular size.

A consequence of equivalent hydration effects *per mass* is the existence of a common “universal” value (heat capacity) for the melting of globular proteins ^1, 2^. However, this heat capacity value is only “universal” when normalized *per gram*, not *per mol* ^1, 2^. When normalized by mass, native heat capacities for individual proteins are essentially the same ^1, 2^. As shown below, equivalence of the heat capacities *per mass* for Proteins and DNA is consistently maintained at the level of individual amino acids and base subunits. Consistent with the proposition that contributions of hydration effects to the absolute heat capacity are essentially equivalent for protein and DNA.

For proteins, it has been reported that, a fundamental relationship exists between mass, transition heat capacity, partial specific volume (PSV), and absolute heat capacity (peak height on DSC thermograms)^4^,

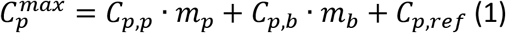

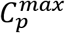, is the maximum calorimetric peak height (measured absolute heat capacity at T_m_) from the thermogram. C_p,p_, transition heat capacity of the protein; m_p_, mass of the protein; C_p,b_, intrinsic heat capacity of the buffer; m_b_, mass of the buffer; and C_p,ref_, the instrumental heat capacity. To account for the volume of the calorimetric cells, buffer and protein mass are expressed as a function of density, ρ_x_, and cell volume, v, where *m*_*x*_=*ρ*_*x*_· *v*. In an ideal solution, the buffer and cell volumes are the same. However, depending on the mass and partial specific volume, *PSV*_*p*_, of the protein, in solution the protein occupies a portion of the calorimetric cell volume. The volume of protein lowers the apparent intrinsic heat capacity of the buffer. To account for this, equation (1) can be rewritten as,

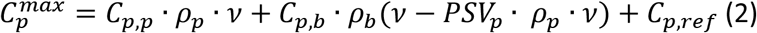

Grouping and factoring the protein mass dependent terms gives equation (3),

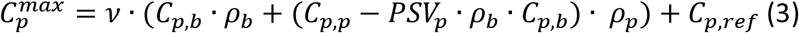

Since all measurements must be made on identical instruments, buffer and instrumental baselines can be directly subtracted, enabling omission of the instrumental term, C_p,ref_, in equation (3). For dilute aqueous solutions, *C*_*p,b*_·*ρ*_*b*_= 1 cal/K·mL and the cell volume, v, is constant for all experiments ^4^. After buffer subtraction, the buffer terms are removed from equation (3) providing,

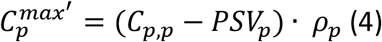

This relationship can be used to determine transition heat capacities for proteins by plotting the maximum thermogram peak heights at 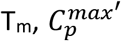, versus mass of protein in the calorimeter ^4^. The same scheme can also be applied to DNA. Since thermograms are not normalized for mass or molecular weight, the same process is applied to determine the observed transition heat capacity of DNA with a slight modification of equation (4),

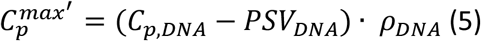

Using equations (4) and (5) the universal transition heat capacities, for proteins and DNA are evaluated.

## Results

Rather than raw C_p_ versus T curves, our analysis employs thermograms after buffer background subtraction and baseline fitting. This process was done to eliminate buffer dependent terms in equation (3). To demonstrate the equivalence of transition heat capacity for proteins, human serum albumin (HSA) and lysozyme were used as examples of typical globular proteins ^2, 4, 16, 20, 21^. An identical analysis was performed to evaluate the transition heat capacity of a DNA hairpin.

To determine transition heat capacities, C_p,p_, for the individual proteins, as a function of mass, thermograms were measured for mass titrations of HSA and lysozyme. C_p,p_ was determined using equation (4). In Fig 1, calorimetric peak height (absolute heat capacity, 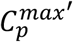, is plotted versus protein concentration (mg/mL). According to equation (4) the slope of the resulting plot equals the transition heat capacity minus the partial specific volume (Slope= C_p,p_ - PSV_p_). Rearrangement of this equation provides C_p,p_, i.e.

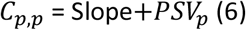

**Figure 1:**
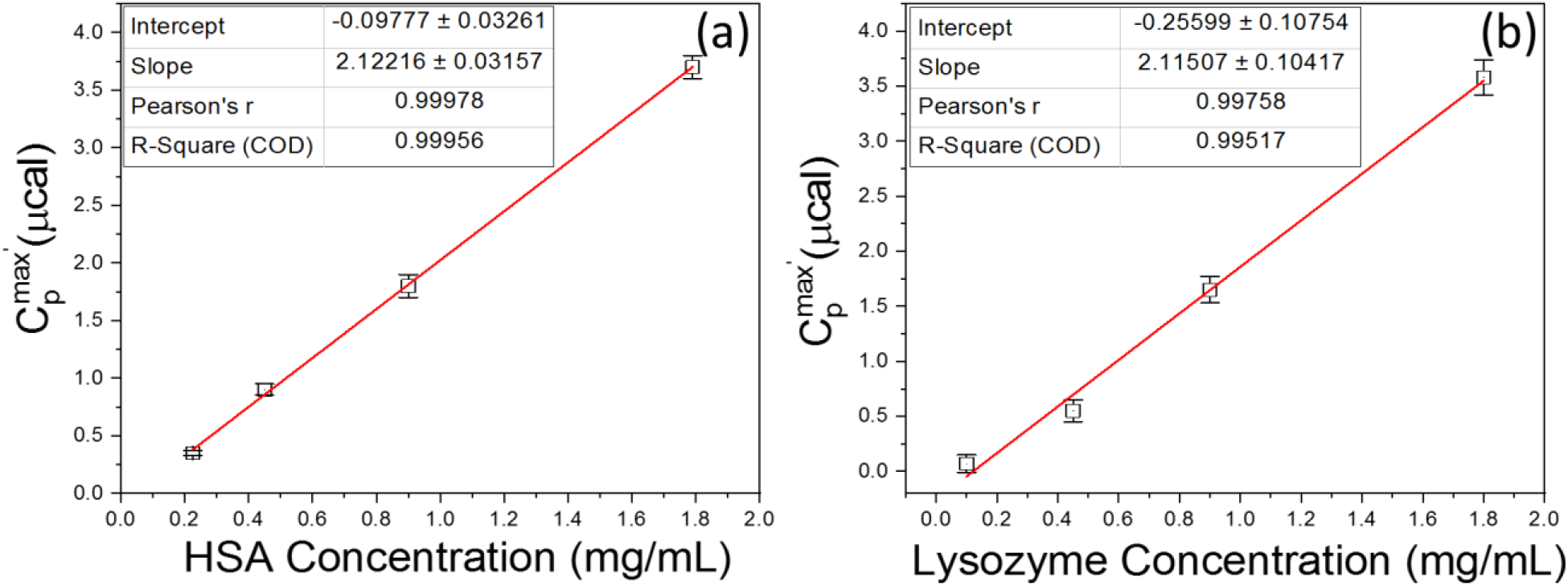
Transition heat capacity curves for HSA (a) and Lysozyme (b) at the T_m_.

Four concentrations (0.23, 0.45, 0.9, and 1.8 mg/mL) for HSA and lysozyme were used to determine C_p,p_. Three measurements were made for each protein solution. Results are shown in Fig 1. Since T_m_ should be independent of the protein concentration, as expected, across the measured concentration range the T_m_ remains unchanged. Measured T_m_ values for HSA and lysozyme were T_m,HSA_ =63.62 ± 0.11°C and T_m,Lys_ =64.77 ± 0.16°C.

PSV values for the proteins taken from the literature were 0.733 mL/mg for HSA and 0.703 mL/mg for lysozyme ^2, 4, 5, 20, 21^. The observed transition heat capacity values for HSA and lysozyme were evaluated, from equation (6), C_p,HSA_= 2.86 ± 0.03 mcal·g^-1^·K^-1^, C_p,Lys_= 2.82 ± 0.13 mcal·g^-1^·K^-1^. To note, results showed a slightly lower overall transition heat capacity and larger variance for lysozyme compared to HSA. Although of little concern, this small difference was likely due to the poor shelf stability of lysozyme solutions, which required preparation of multiple stock solutions and more precise timing of measurements. Even with these factors effecting the lysozyme samples, nearly exact C_p,p_ values were obtained, supporting the assertation that, normalized per mass, the transition heat capacity is essentially the same for these globular proteins.

The DNA was a 20-base single strand oligomer designed to form an intramolecular stem-loop “hairpin” structure; with six base pairs in the duplex stem and four-base single strand loop on one end. The stability of the six base pair sequence was designed to have a melting temperature above 90°C; T_m,DNA_ =94.34 ± 0.06°C. Again, as expected, this T_m_ was independent of DNA concentration, with no evidence of intermolecular duplex formation.

DSC thermograms were collected for DNA at concentrations from 0.02 mg/mL to 2.50 mg/mL. Experiments were performed in triplicate. Maximum peak heights on DSC thermograms, ^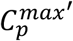^ values, were determined for each DNA concentration, resulting in the curve shown in Fig 2. The excellent linearity of the plot of 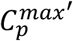 versus DNA concentration (R^2^ = 0.999) confirms the linear dependance on mass for the DNA sample.

**Figure 2:**
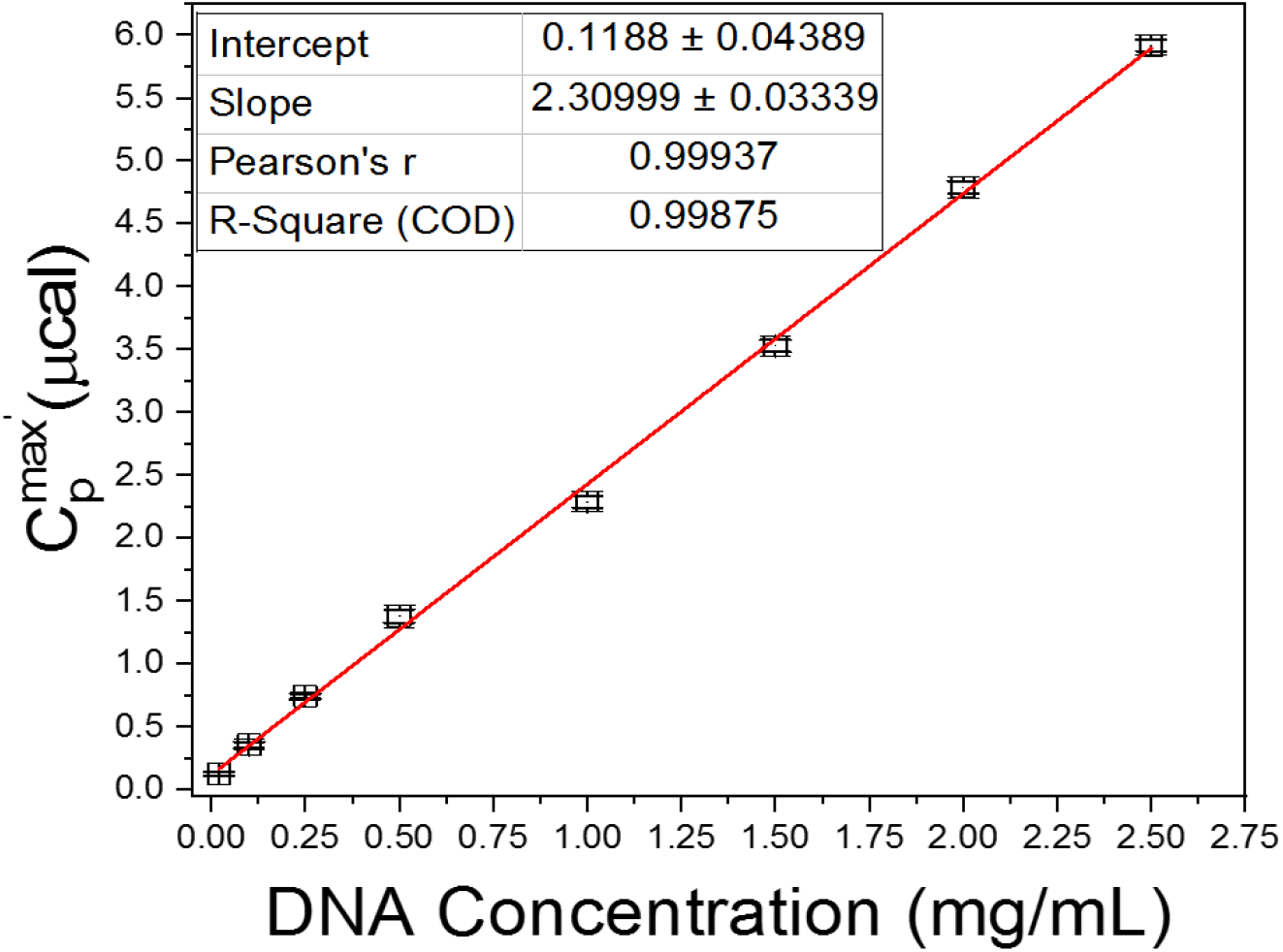
DNA hairpin transition heat capacity curve at the T_m_.

In an analogous manner to the proteins above, equation (6) was used to evaluate the observed transition heat capacity for the DNA hairpin. The literature value for short DNA oligomers PSV_DNA_=0.55 mL/g, was employed ^22^. With this PSV, the transition heat capacity for the DNA hairpin was determined to be C_p,DNA_= 2.86 ± 0.03 mcal·g^-1^·K^-1^.

Thus, C_p,DNA_ (2.86) is precisely the same as C_p,HSA_ (2.86) and nearly identical to C_p,Lys_ (2.82). Although only demonstrated for these three molecules, these results are consistent with “universal” values that have been evaluated for many proteins at 25°C. Although for only a single molecule examined here, the agreement of C_p,DNA_ with those of the proteins, suggests the potential equivalence for transition heat capacity values of proteins and DNA. Although attractive, the generality of this proposition must be confirmed for additional DNAs.

The above results can be used to evaluate the transition heat capacity of individual molecules, C_p,DNA_ = 3.63 × 10^−20^ mcal·molecule^-1^·K^-1^ for DNA and C_p,HSA_ = 3.16 × 10^−19^ mcal·molecule^-1^·K^-1^ for HSA. This indicates that each HSA molecule has 8.7 times as much heat capacity as a single DNA molecule. Coincidently, the ratio of molecular weights of HSA to DNA is also 8.7. Thus, at a given mass in a sample, there will be 8.7 times more DNA molecules than HSA molecules. Likewise, for lysozyme C_p,Lys_ = 6.79 × 10^−20^ mcal·molecule^-1^·K^-1^. Each lysozyme has 1.9 times as much heat capacity as a single DNA molecule, where again the ratio of molecular weights of lysozyme to DNA is also 1.9.

Examining the heat capacity per amino acid or base residue for each molecule, C_p,HSA_ = 5.40 × 10^−22^ mcal·residue^-1^·K^-1^, C_p,Lys_ = 5.26 × 10^−22^ mcal·residue^-1^·K^-1^ and C_p,DNA_ = 1.81 × 10^−21^ mcal·base^-1^·K^-1^. Thus, the heat capacity of each DNA base is 3.36 and 3.45 times greater than an amino acid residue for HSA or lysozyme, respectively. Coincidently, this corresponds to a factor of 3.36 greater for the average molecular weight of each base of the DNA hairpin (382.1 g/mol) compared to each amino acid of HSA (113.7g/mol). A similar factor 3.45 exists for the ratio of the molecular weight of each DNA base to lysozyme amino acid (110.0 g/mol). This observation suggests the heat capacity of proteins and DNA corresponds to the mass of individual bases or amino acid residue. Interestingly, this is entirely consistent with the proposition that molecular size is directly coupled to hydration enthalpy ^13, 15^. Said another way, per residue, since DNA bases are approximately three times larger than amino acid residues their hydration enthalpies are likewise three times greater.

Observation of identical heat capacities for DNA and proteins, C_p,p_ = C_p,DNA_, is intriguing and potentially implies a common origin for protein and DNA global structural stability due to hydration effects.

While water and hydration effects are the most likely cause for this phenomenon, “hydration” is a simplistic explanation. As mentioned, water has multiple effects on protein stability from the native hydration shell to hydration of hydrophilic and hydrophobic groups. Within these effects there is likely enthalpy/entropy compensation that, *per mass*, is normalized for the specifics of small DNA oligos and large protein molecules. The exact nature and magnitude of hydration effects between large and small molecules is currently the subject of investigation.

There are practical applications for these results. For example, because of the equivalence of the transition heat capacity of proteins and DNA, the unknown mass of a protein can be quantitatively evaluated solely from the measured calorimetric peak height of the protein, when compared to the peak height for a known amount of DNA of the same mass.

Note: The transition heat capacity values we evaluated for lysozyme and HSA differ from those previously reported in the literature for the same proteins ^2, 11^. The origin for this difference arises from our use of the transition heat capacity at T_m_ compared to the native heat capacity evaluated at 25°C reported in the literature. Nevertheless, within the same data group, i.e. same instrument, same baseline parameters, same data treatment, our data evaluated at T_m_ is internally consistent and again demonstrates the universal heat capacity for globular proteins at any temperature. This uncertainty underscores the need for internal standardization prior to using DSC to evaluate masses of unknown proteins.

## MATERIALS AND METHODS

### Chemicals and Reagents

Standard buffer for all experiments contained 150 mM NaCl, 10 mM potassium phosphate, 15 mM sodium citrate adjusted to pH = 7.4 with hydrochloric acid. Pure Proteins: Human Serum Albumin (HSA) (≥ 99% pure, Lot number: SLBT8667) and Lysozyme (recombinant, expressed in rice, Lot number: SLCH2681). The above proteins were prepared in standard buffer and stored at 4°C for at least 24 hours before use. Concentrations of HSA and lysozyme were confirmed spectrophotometrically at A_280_^20, 23^.

### DNA

The DNA was a 20-base pair hairpin purchased from IDT and received following a standard desalting routine. The DNA standard sequence is 5’ -CGG GCG CGT TTT CGC GCC CG-3’. Lyophilized DNA was resuspended in standard buffer and stored at 4°C. DNA concentration matched the manufacturer specification as determined spectrophotometrically at A_260_.

### DSC Measurements

DSC melting experiments were made using a CSC differential scanning microcalorimeter (now T.A. instruments, New Castle, DE). For DSC melting experiments, the sample heating rate was approximately 1 °C/min while monitoring changes in the excess heat (microcalories) of the sample versus temperature ^5, 21, 24^. Sample volumes for DSC melting experiments were 0.5 mL.

### Data Reduction and Analysis

Thermograms of proteins were displayed in primary form as plots of microcalories (*μcal*) versus temperature. Baselines of the raw *ΔC*_*P*_ or *μcal* versus temperature thermograms were determined using a four-point polynomial fit, over the temperature range of the transition, consistent with our previous methods (citations). For all protein mixtures the buffer baseline was then subtracted from the raw curves, producing baseline corrected thermograms used for further comparisons and analysis.

Linear fits of peak height versus protein/DNA concentration were performed in triplicate and fit using Origin(Pro), Version 2021b, Origin Corporation, Northampton, MA.

## Supplemental Material

**Figure S1:**
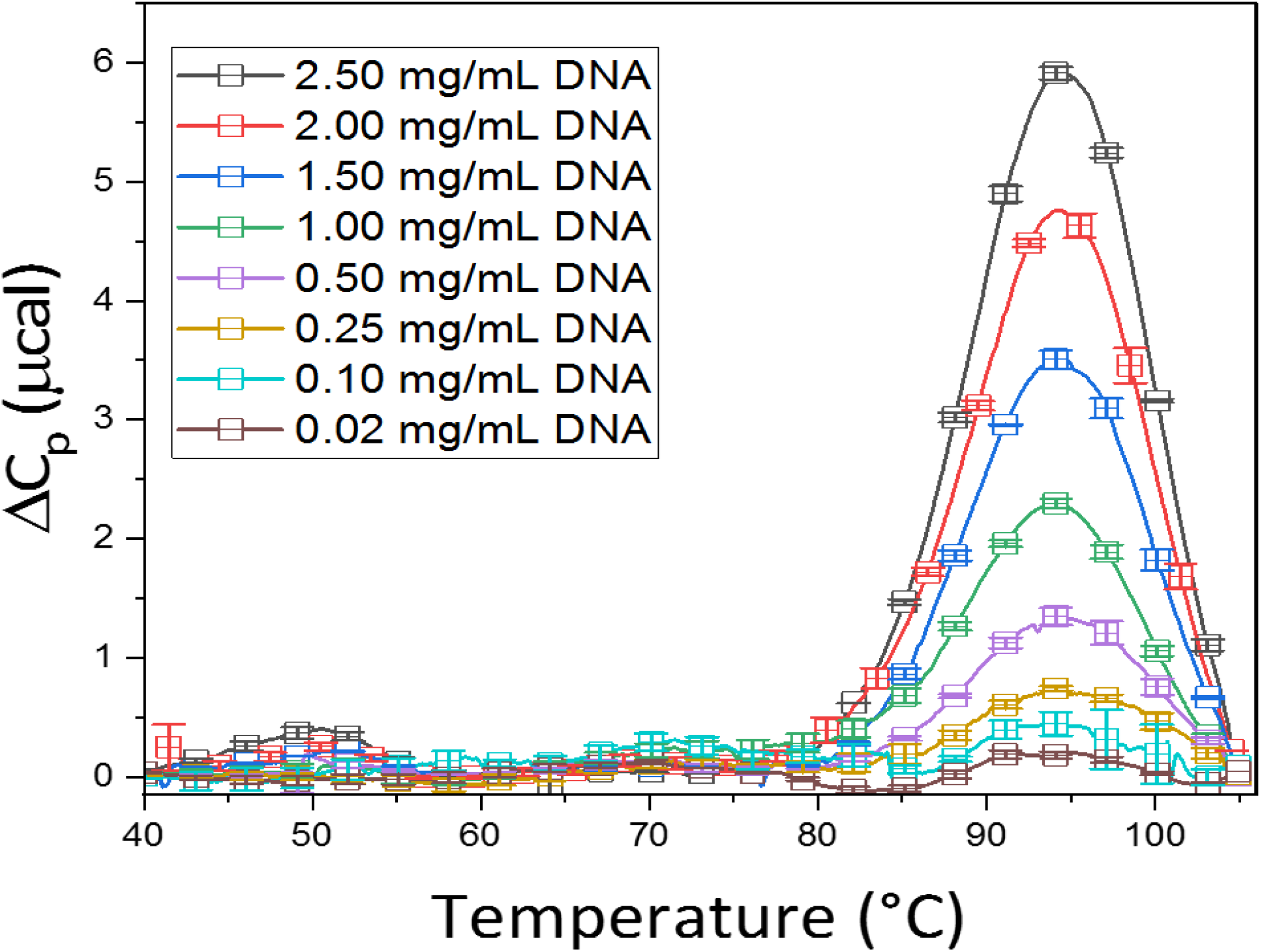
Average thermograms for DNA hairpin from 2.5-0.02 mg/mL.

**Figure S2:**
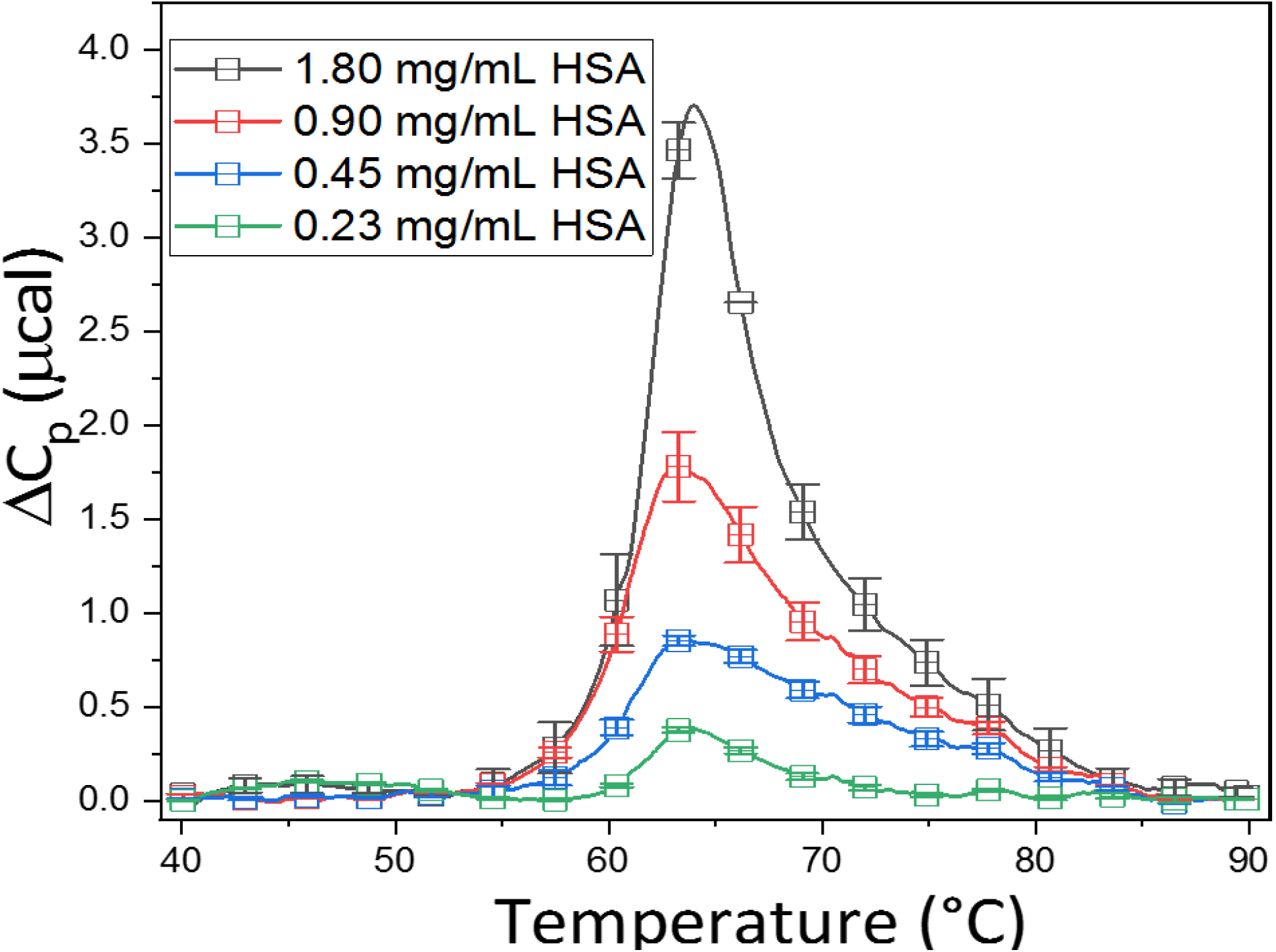
Average thermograms for human serum albumin (HSA) from 1.8-0.23 mg/mL.

**Figure S3:**
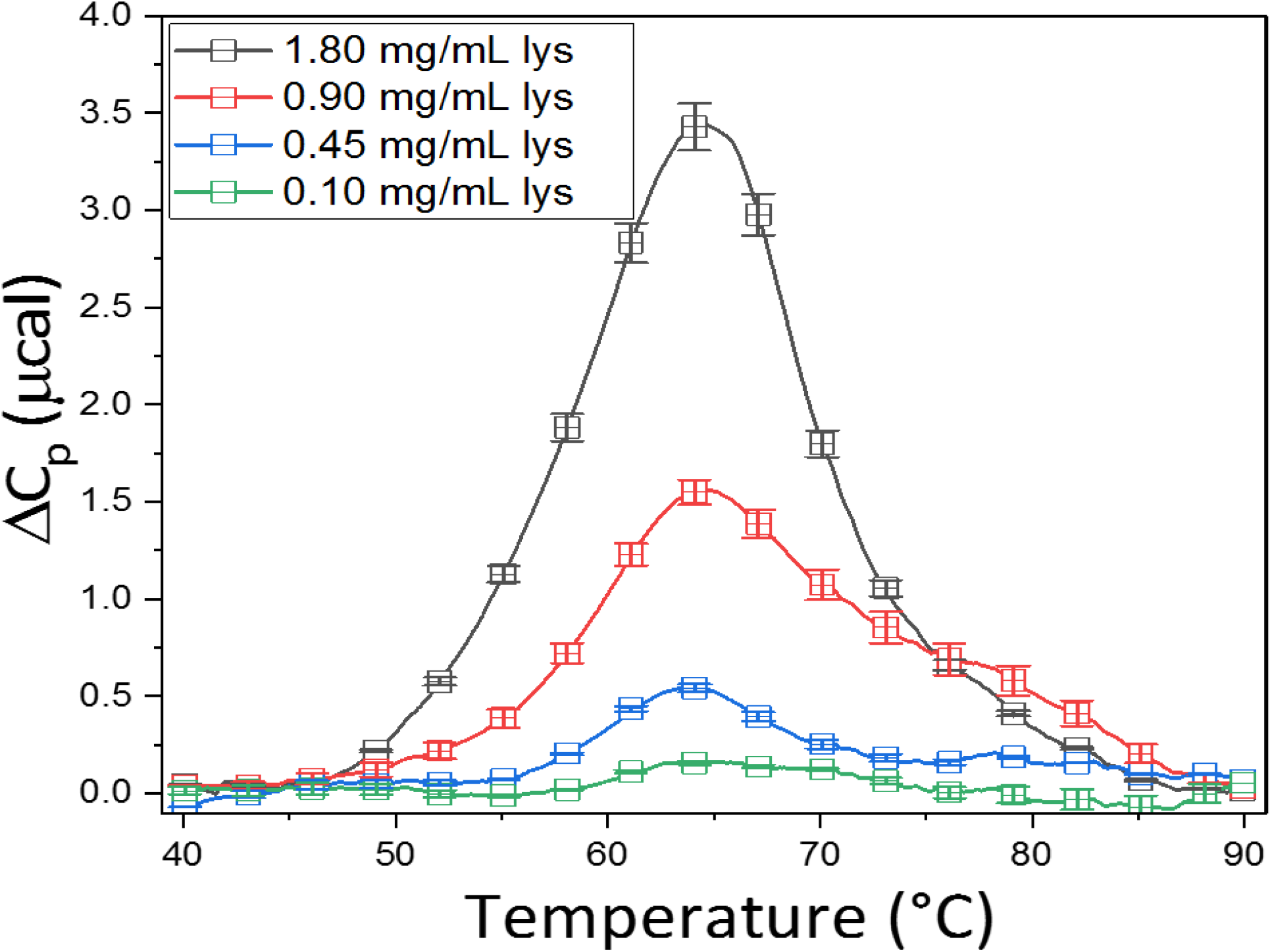
Average thermograms for lysozyme (lys) from 1.8-0.1 mg/mL.

